# Gonadal Transcriptome Differences Between the Sexes in Wild Caught Southern Flounder (*Paralichthys lethostigma)* During a Critical Development Period

**DOI:** 10.1101/2025.10.21.683694

**Authors:** Sydney Harned, Jamie L. Mankiewicz, Ethan Smith, Russell J. Borski, Rafael F. Guerrero, Martha Burford Reiskind, John Godwin

**Author notes:** **Corresponding author:** Sydney P. Harned, M.S.

## Abstract

Flounder in the genus *Paralichthys* exhibit a unique sex determination system that combines genotypic and temperature-dependent mechanisms. In southern flounder (*Paralichthys lethostigma*), XY individuals develop as males, while XX individuals’ sex is influenced by juvenile developmental temperatures. Little is known about the genes involved in this process. This study uses pooled RNA sequencing to create a de novo gonadal transcriptome and identify genes related to sex determination and differentiation in wild-caught juvenile southern flounder from two North Carolina locations with differing sex ratios. The transcriptome assembly revealed 68,331 uniquely annotated genes with high functional specificity. Nine genes were linked to sex determination, including five novel to southern flounder and associated with epigenetic mechanisms like m6a methylation and alternative splicing, suggesting epigenetic influence in male development. Additionally, genes in the TGF-β signaling pathway were differentially expressed, supporting this pathway’s role in sex differentiation. This study provides a valuable transcriptome resource for future research on southern flounder.

## Introduction

The remarkable plasticity of sex determining systems in fish presents a distinctive challenge in identifying sex-determining genes and pathways [1–2]. Teleost fish demonstrate diverse sex-determining systems, and the evolutionary lability of these systems has led to the independent evolution of sex-determining mechanisms even among closely related species [2–3]. Many fish species undergo environmental sex determination [4], where external factors such as temperature drive the sex determination and differentiation process [5–7]. As environmental factors continue to shift rapidly due to climate change, species that undergo environmental sex determination are at risk of highly skewed sex ratios and ultimately population decline or extinction. As such, there is an urgent need to understand the extent of environmental sex determination in wild populations of fish exposed to rapidly changing environments.

In southern flounder, *Paralichthys lethostigma*, a combination of genetic factors and environmental temperature influence sex determination. Previous studies suggest this species has a heterogametic XX/XY sex determination system (GSD), where genetic or epigenetic factors trigger either male (XY) or female (XX) development [7–10]. However, environmental cues can override this genetic system. Specifically, during the early post-settlement juvenile period (approximately 35-65 mm total length), exposure to rearing temperatures either higher or lower than 23°C in North Carolina southern flounder or above approximately 18°C for Texas southern flounder significantly increases the proportions of phenotypic males (>90% at 28-30°C in [7] and [11] respectively). Despite this established pattern of temperature-dependent sex determination (TSD) in southern flounder and some studies of its molecular correlates [8–9], the genetic factors governing sex determination and TSD in this species remain largely unknown, and the master sex-determining gene has yet to be identified.

Sex reversal of genetically XX female flounder can have critical implications for this important fishery resource. The southern flounder fishery has been declining for several decades [12–13], potentially because of increased temperatures in nursery habitats [14], and a recent population modeling study predicts a continued decline with increasing male-bias [15]. As females are 2-3 times larger in this species, male southern flounder often fall short of the minimum size requirement for harvest (350 mm in North Carolina; [12]), resulting in a predominance of females in recreational and commercial landings. This female-biased harvesting pattern poses a considerable threat to the sustainability of the southern flounder fishery. A heavily skewed sex ratio leads to fewer reproducing females and a low spawning stock biomass. Southern flounder are also important predators in their shallow estuarine ecosystems, and a decline in abundance can cause an imbalance in lower trophic level species [16–17]. Therefore, there is a need to understand the prevalence of sex reversal and TSD in southern flounder. To address this, it is valuable to identify the genetic factors governing sex determination.

Several early markers of sex determination have been identified and can be used to determine phenotypic sex in southern flounder. These markers include gonadal aromatase (*cyp19a1a*) and forkhead-box protein L2 (*foxl2*) in the female development pathway, and the anti-mullerian hormone (*amh* or Müllerian inhibiting substance*, mis*) in the male development pathway [9, 11, 18]. These genes have known roles in the sex-determining pathway in several teleost species [19–23] While these markers are useful in understanding and identifying the phenotypic sex of southern flounder during early juvenile development [8, 24], the possible existence and identity of a master sex-determining gene remains elusive. Importantly, a Y-specific variant of the *amh* gene, *amhy*, was recently identified as a likely master sex-determining gene in the olive flounder, *Paralichthys olivaceus* [22]. Whether this *amh* gene variant is the master sex determining gene or other unknown factors are crucial to sex determination in southern flounder remains unclear. In recent years, members of the TGF-β signaling pathway have been consistently identified as master sex-determining genes in a range of species, including fishes [25–28]. Of 114 fish species where the master sex-determining gene has been identified, 72 of these belong to the TGF-β signaling pathway, suggesting this pathway plays a key role in sex determination [26]. *Amh*, the master-sex determining gene in the olive flounder, is a member of the TGF-β family; however, the role of TGF-β genes in southern flounder sex determination has not been investigated beyond use as a molecular biomarker of sex [9, 24].

The goal of this study is to advance our understanding of sex determination in southern flounder via transcriptomics and measures of differential gene expression at a more genomic scale through RNA sequencing. We assembled a transcriptome for southern flounder and described gene expression differences during critical developmental stages between wild-caught males and females. Our investigation focused on identifying gene expression differences between sexes in gonad tissue of the early sex determination and differentiation period for wild-caught juvenile southern flounder sourced from two distinct North Carolina estuaries characterized by geographic and temporal variation and different presumed rates of TSD – one with approximately even sex ratio and the other exhibiting heavily male-biased juvenile sex ratios [24]. We found four well-established sex-determining genes. Additionally, we identified five novel candidate genes that may play crucial regulatory roles within the male sex determination and differentiation pathway, and other genes involved in the TGF-β pathway that may also be involved in sex determination. Identifying these genes offers a chance to better understand sex determination and temperature effects in southern flounder, and to develop new biomarkers for early juvenile phenotypic sexing.

## Methods

### Sampling and RNA Extraction

We obtained juvenile samples from two estuaries: Pamlico Sound, characterized by an approximately 50:50 juvenile sex ratio, and Mill Creek, where juvenile sex ratios are skewed up to 94% male (Fig. 1; [24]). The North Carolina Division of Marine Fisheries collected juvenile southern flounder from North Carolina upper estuaries through their routine juvenile monitoring Program 120 (P120) trawl survey using an otter trawl towed for 1 minute at 1 m/s. We also used additional samples collected as described in [24].

**Fig. 1.**
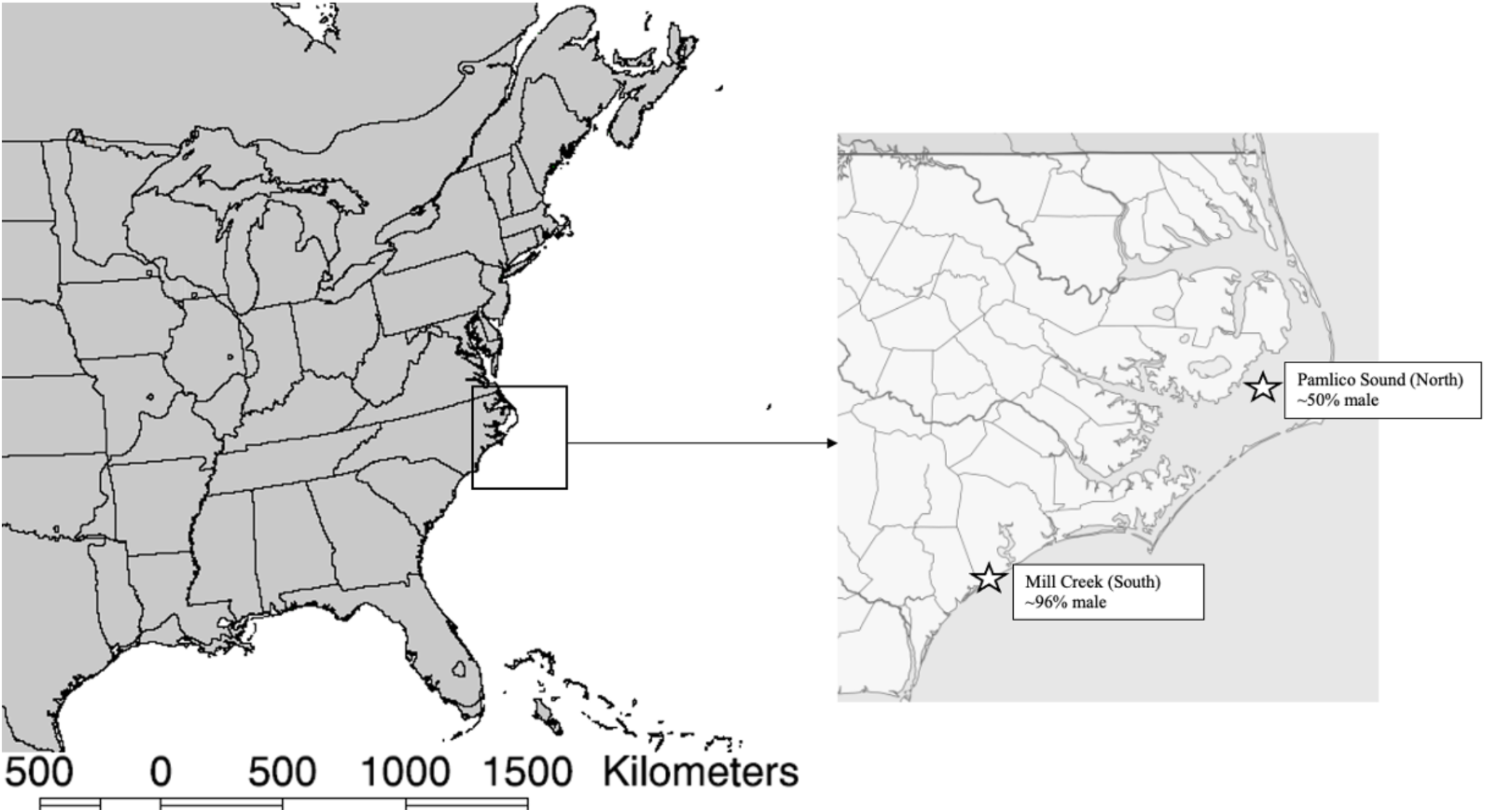
Southern flounder sampling locations in North Carolina. Proportion of males based on Honeycutt et al. 2019.

For the subsequent RNA extraction process, we extracted total RNA from gonadal tissue, quantified and DNAse treated it as previously described [24]. RNA was quantified using absorbance OD 260:280 ratio measured by a nanodrop spectrophotometer (Nanodrop-1000, ThermoFisher Scientific, Waltham, MA) and quality checked via a BioAnalyzer at the North Carolina State University Genomic Sciences Laboratory (Raleigh, NC).

### RNA Pooling and Sequencing

We pooled RNA from individual flounder samples based on phenotypic sex (determined by molecular biomarkers, as outlined in [24]), total length in 10 mm increments (50-59mm, 60-69mm, and 70-79mm), and sampling location (Pamlico Sound - northern population; Mill Creek - southern population). We diluted each RNA sample to 100 ng/ul per individual and combined these individual samples into pools representing 10 individual fish, resulting in a total pool of 1 ug RNA. Two pools, representing southern males of 50-60 mm TL and 60-70 mm TL, contained 12 and 9 individuals, respectively, due to constraints in RNA availability (Table 1). We then checked the pools for quality and concentration via BioAnalyzer and carried out sequencing via Illumina Hi-Seq at the NC State Genomic Sciences Laboratory.

**Table 1.** Southern flounder sample distribution and pooling.

| **Pool Name** | **Location** | **Sex** | **Size (TL)** | **N** |
| --- | --- | --- | --- | --- |
| Northern_Female_60 | North | F | 60-69 mm | 10 |
| Northern_Female_70 | North | F | 70-79 mm | 10 |
| Northern_Male_50 | North | M | 50-59 mm | 10 |
| Northern_Male_60 | North | M | 60-69 mm | 10 |
| Northern_Male_70 | North | M | 70-79 mm | 10 |
| Southern_Male_50 | South | M | 50-59 mm | 12 |
| Southern_Male_60 | South | M | 60-69 mm | 9 |
| Southern_Male_70 | South | M | 70-79 mm | 10 |

### De Novo Transcriptome Assembly and Quantification

We employed the *de novo* transcriptome assembly software Trinity v2.15.1 to assemble paired-end Illumina RNA-seq reads. Trinity is a specialized software package designed for the robust construction of transcriptomes from RNAseq data [29]. We ran Trinity with a minimum contig length of 200 bp and a normalized maximum read coverage of 200.

### Assembly Quality Control

To assess accuracy and reliability of the *de novo* transcriptome assembly, we employed the rnaQUAST v2.2.3 software package [30], implemented in a Python v3.9.5 environment. We ran rnaQUAST with a minimum alignment threshold of 50 and lower and upper thresholds set at 50% and 95%, respectively. To assess transcriptome completeness, we ran the widely used software tool BUSCO (Benchmarking Universal Single-Copy Orthologs; [31]) using the default settings. BUSCO evaluates the completeness of the assembled transcriptome by comparing it to a set of universal single-copy orthologs. The dual approach using rnaQUAST and BUSCO enhances the robustness of the quality control process, offering a comprehensive assessment of the assembly.

### Transcript Quantification

We estimated transcript abundance with the ‘align_and_estimate_abundance.pl’ utility script from Trinity [32] with the ‘alignment-free’ method, employing the mapping-based mode of the salmon tool v1.10.0 [32–33] for transcript quantification. The parameters passed to the ‘align_and_estimate_abundance.pl’ script were as follows: --seqType fq, --est_method salmon, -- trinity_mode, --prep_reference, --salmon_idx_type quasi (default). The ‘salmon_idx_type quasi’ parameter designates the quasi-indexing type as the default setting. Subsequently, we constructed transcript and gene expression matrices using Trinity’s ‘abundance_estimates_to_matrix.pl’ utility script [32]. This step facilitates the organization and compilation of transcript abundance data, providing a basis for downstream analyses.

### Functional Annotation of Transcriptome Assembly

To functionally annotate the Trinity *de novo* transcriptome assembly, we utilized the Trinotate annotation suite v4.0.0 [34], an open-source tool designed for gene identification and functional annotation of transcripts. Trinotate derived annotations from BLASTx and BLASTp top hits from the Swiss-Prot and Uniref90 databases, SignalP, TmHMM, Pfam domain, eggnog, Gene Ontology (GO) from BLAST, and GO from Pfam.

The annotation process involved several steps. Initially, we created an SQLite database to store annotation data by executing the Trinotate software package with the ‘--create’ flag.

Subsequently, TransDecoder was employed to predict likely coding regions within the transcripts from the assembly (Haas, BJ. https://github.com/TransDecoder/TransDecoder). Following the generation of a file containing predicted coding regions translated in FASTA format by TransDecoder, the Trinotate SQLite database was initialized with the sequence data using the ‘--init’ flag. Finally, we generated annotations for each transcript with Trinotate.

To achieve comprehensive homology searches, we utilized the Swiss-Prot and Pfam databases alongside BLAST+ v2.11.0 and HMMER v3.3.2 [35–38]. Infernal v1.1.4 was also incorporated during the Trinotate run for non-coding RNA identification [39]. After completion, we extracted the annotation information to a tab-separated values (tsv) file by running Trinotate with the ‘--report’ flag. The resulting file contains information regarding the functional annotation of the *de novo* transcriptome assembly, providing insights into the homology and function of identified transcripts.

### Identification of Differentially Expressed Genes in Males vs. Females

We identified differentially expressed genes (DEGs) between male and female southern flounder using the Bioconductor package edgeR in R and gene count data produced in Salmon for each pool [40]. We used a fixed dispersion value of 0.1 in edgeR to account for a lack of biological replicates. We performed two separate comparisons: northern females vs. northern males and northern females vs. southern males. The southern location is likely to have a high rate of sex reversal, and southern females are very rare (<5%, [24]) and thus not available for this study.

Therefore, we use northern females as the comparison for both northern and southern males. In each comparison, we restricted analysis to size classes where both male and female data were available, specifically individuals ranging from 60-69 mm TL and 70-79 mm TL. We set criteria for determining significant DEGs at a false discovery rate (FDR) adjusted p-value of <0.01 and a log2 fold change (FC) of ≥2 or ≤-2. We considered transcripts meeting these criteria as significantly differentially expressed.

To identify genes potentially involved with sex determination and differentiation, we first searched the list of differentially expressed genes for those previously identified as sex-determination genes in southern flounder and closely related species. These genes included *mis* (or *amh), dmrt1, cyp19a1a,* and *foxl2.* Then, to search for sex-related genes that have not been characterized in southern flounder, we extracted transcripts with Gene Ontology (GO) terms related to sex determination and differentiation. The relevant GO terms included: GO:0007530 biological_process_sex determination, GO:0046661 biological_process_male sex differentiation, GO:0046660 biological_process_female sex differentiation, GO:0030238 biological_process_male sex determination, GO:0030540 biological_process_female genitalia development, and GO:0007548 biological_process_sex differentiation. We classified transcripts as candidate sex genes if they met the following criteria: 1) LogFC of >2 or <-2 for up- or down-regulated transcripts, respectively; 2) FDR <0.01; 3) consistent protein homology across databases (BlastX, BlastP, PFAM); and 4) for males, consistent differential expression relative to northern females in both southern and northern pools. These criteria ensure a stringent selection of candidate sex genes, providing a foundation for further exploration of their roles in sex determination and differentiation in southern flounder. To assess the role of TGF-β genes in southern flounder sex determination and differentiation, we also conducted a search for key TGF-β genes associated with sex in other fish species (Table 1 in [27]).

## Results

### De novo transcriptome assembly

The assembly, built from 1,650,725,544 100-bp paired-end reads from all pools, yielded a total of 485,040 transcripts. Of these, 284,478 transcripts had a length of >500 base pairs, and 169,390 transcripts had a length of >1000 base pairs, signifying a substantial representation of longer transcripts (Table 2). The Transcript N50, indicating the length at which half of the assembled bases are incorporated into transcripts of that length or longer, was 2218 base pairs. The average length of the transcripts was 1198.73 base pairs. The assembly also predicted the presence of 151,195 genes, representing potential coding regions within the transcriptome.

**Table 2.** Trinity assembly metrics for the southern flounder *de novo* transcriptome.

|  |  |
| --- | --- |
| Transcripts | 485040 |
| Transcripts > 500 bp | 284478 |
| Transcripts > 1000 bp | 169390 |
| Complete | 100.0 |
| Partial | 0.0 |
| Predicted genes | 151195 |

### 3.2 Assembly Quality Control

Both RNAQuast and BUSCO determined 100% completeness of the *de novo* transcriptome. BUSCO yielded 255 complete hits, and all expected single-copy orthologous genes were identified in the transcriptome assembly. Of the complete hits, 43 (16.9%) were identified as single copies, while 212 (83.1%) were classified as duplicated copies. The presence of duplicate copies probably indicates allelic variation within the total sample. Since all pools were utilized in constructing the transcriptome, this observation aligns with the expectation of capturing genetic diversity, including potential allelic differences, within the wild-caught samples from this southern flounder population. The absence of fragmented or missing BUSCO hits further underscores the robustness and completeness of the *de novo* transcriptome assembly.

### Transcriptome annotation

A total of 303,233 transcripts were subjected to Trinotate annotation using reference genomes from a range of species; of these, 95,991 transcripts received at least one annotation. These annotations provide information about the putative functions of the identified transcripts. A total of 68,340 genes were identified in Trinotate. Among these genes, 68,331 were uniquely annotated, indicating a high degree of specificity in assigning functional information to these genes. Additionally, nine genes had duplicate annotations, possibly suggesting instances where multiple transcripts are associated with the same gene due to alternative splicing or other factors. The nine duplicated transcripts were kept in the dataset.

### Differential gene expression analysis between males and females

We assessed the set of 68,331 uniquely annotated genes for differential expression between males and females. In the comparison between northern males and females, we found a total of 3082 genes to be significantly upregulated in males, with 2912 of these genes annotated in Trinotate (Table S1). Conversely, 739 genes were significantly upregulated in females, and 714 of these genes received annotations (Table S2). By applying filtering based on sex-related Gene Ontology (GO) terms, we identified seven upregulated sex-related genes in males, and two in females.

In comparison between southern males and northern females, 9241 genes were significantly upregulated in males, with 8794 of these genes annotated in Trinotate (Table S3). For females, 1652 genes were significantly upregulated, and 1590 of them were annotated in Trinotate (Table S4). Similar to the Northern Pamlico comparison, we identified seven upregulated sex-related genes in males, and two in females.

The seven sex-related genes upregulated in males and the two in females were consistent between the two comparison groups (Table 3). The seven male-associated genes were homologous to *Protein Groucho*, *Splicing Factor 1*, *Cullin*, RNA-*Binding protein Spenito*, *N6-Adenosine-Methyltransferase*, *mullerian-inhibiting factor,* and *dmrt1*. The two female genes were homologous to *cyp19a1a* and *foxl2*. These consistent findings across different comparison groups strengthen the confidence in the identification of sex-related genes that may play crucial roles in the sex determination and differentiation processes in southern flounder.

**Table 3.**
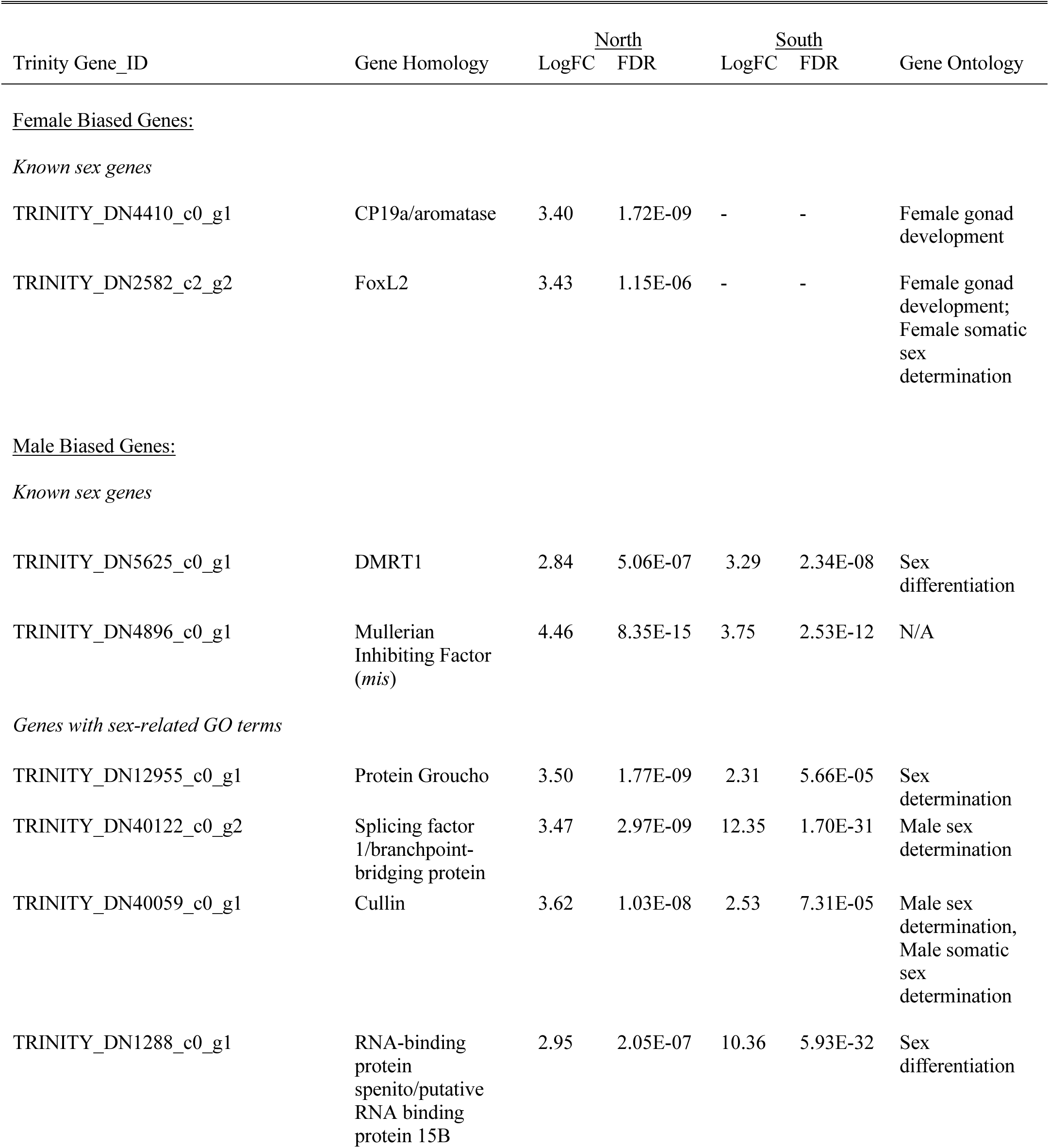
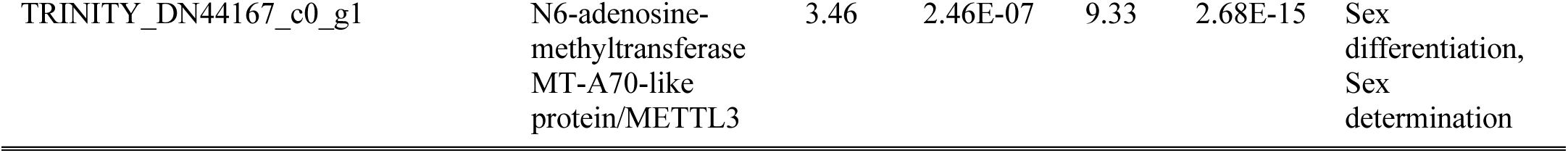
Significantly differentially expressed genes with potential sex-related function between the gonads of male and female southern flounder transcriptomes.

Several genes associated with the TGF-β gene family were significantly differentially expressed in at least one male to female comparison (Table S5). One of these, transforming growth factor-beta receptor associated protein 1, was elevated in both northern and southern males relative to northern females. Other differentially expressed transcripts associated with TGF-β signaling included those implicated in bone morphogenetic protein signaling and latent TGF-β binding protein. There was some indication of differential expression of SMAD protein mRNAs, which participate in TGF-β signaling and have been proposed to play a role in TSD in Japanese flounder [26], being differentially expressed between southern males and northern females (Table S3, row 2045). However, this SMAD4 transcript was not significantly differentially expressed in comparisons of northern males to northern females (Table S1, row 6674). Lastly, there was significantly higher expression of transient receptor potential (TRP) cation channel transcripts in females, most prominently with TRPM5.

## Discussion

### Differential expression of known sex markers

Our study found differentially expressed, previously documented markers of female sex determination and differentiation in southern flounder, *foxl2* and *cyp19a1a*. We found these genes upregulated in females, consistent with previous studies of sex markers in southern flounder and closely related species; both *foxl2* and *cyp19a1a* increase in expression and remain upregulated during female sex differentiation in both olive and southern flounder [8–9, 41–43]. These genes work together in ovarian differentiation: *cyp19a1a* encodes the gonadal form of the aromatase protein, a critical component of estrogen biosynthesis, while *foxl2* promotes the transcription of *cyp19a1a* [9, 18, 41]. As both markers have been used to phenotypically sex southern flounder via qPCR [24], our findings further validate this method by confirming upregulation of these markers via a different method of quantifying gene expression (qPCR vs. RNAseq).

We identified two known sex markers as upregulated in males, *dmrt1* and *mis*, consistent with previous findings in southern and olive flounder. *Dmrt1* is an early marker of male differentiation, and we found it upregulated in both northern and southern males relative to females. *Dmrt1* inhibits expression of *cyp19a1a* in the male pathway and has been identified as the master sex-determining gene in other teleost species [19,21,44]. The upregulation of this gene in males supports previous findings of its role in male sex determination in *Paralichthyidae* [45]. The *mis* gene, synonymous with and more commonly referred to as *amh* (encodes Anti-Mullerian hormone), is associated with male sex-determination in southern flounder [9] and has been identified as the master sex determining gene across 33 species in 12 genera [27].

Importantly, a variant termed *amhy* was recently identified as the master sex-determining gene in the congeneric olive flounder (*P. olivaceus*; [22]). These findings support previous studies that suggest that these two genes play a pivotal role in sex determination in southern flounder and closely related species. As noted for *cyp19a1a* and *foxl2,* these findings for provide additional validation of qPCR results as well as serving as a ‘positive control’ of sorts for the RNAseq-based comparisons presented here.

Genes in the TGF-β signaling pathway, which includes *mis/amh*, are particularly prominent in sex determination and differentiation across species, including teleost fish [25–27]. Our approach identified several genes from the TGF-β signaling pathway in addition to *mis/amh* that were significantly differentially expressed in at least one comparison (Table S5). Other differentially expressed transcripts associated with TGF-β signaling included those implicated in bone morphogenetic protein signaling and latent TGF-β binding protein. There was also an indication of differential expression of SMAD protein mRNAs, which encode proteins that participate in TGF-β signaling and have been proposed to play a role in TSD in Japanese flounder [26].

SMAD4 was differentially expressed only between southern males and northern females and not in comparisons of northern males to northern females. So, this result should be interpreted with caution. Further studies aimed at exploring the role of TGF-β genes in sex differentiation and determination of southern flounder and other paralichthyids is likely to be fruitful [26].

### Identification of novel putative sex markers

Two genes putatively involved in sex differentiation demonstrated homology to those involved in N6-methyladenosine (m6a) methylation machinery: *RNA protein binding spenito/putative RNA binding protein 15B* and *N6-adenosine-methyltransferase MT-A70-like protein/METTL3*. The involvement of m6a methylation in sex differentiation was observed in various species, including the closely related olive flounder, *P. olivaceus* [46]. Wang et al. found that 60% of differentially expressed genes (DEGs) between olive flounder ovaries and testes exhibited an inverse correlation between expression and m6a methylation, suggesting a role for m6a methylation in the epigenetic regulation of sex pathways [46]. Another upregulated gene in males, *Cullin*, also plays a role in m6a methylation. *Cullin* is associated with the E3 ubiquitin ligase complex, which contributes to the stabilization of the m6a methylation machinery. In olive flounder gonads, ubiquitin ligases containing *Cullin* were enriched by m6a modifications [46]. A previous study identified Ubiquitin ligases as regulators of sex determination in the nematode *Caenorhabditis elegans* [47]. These results suggest that m6a methylation may play a role in epigenetic regulation of the male sex determining pathway in southern flounder.

The identification of two upregulated male genes homologous to alternative splicing factors suggests a role for alternative splicing events in male sex determination. One of these genes shares homology with the *RNA-binding protein Spenito*, a well-known sex-determining gene in the fruit fly *Drosophila melanogaster* that controls the splicing of female development genes. Loss of *Spenito* activity in *Drosophila melanogaster* was associated with female-to-male sex reversal [48]. In addition to *Spenito*, we found another upregulated alternative splicing gene in male southern flounder, broadly denoted as “*splicing factor 1*”. While the specific identity of this splicing factor requires further investigation, its upregulation suggests involvement in splicing events that regulate male sex determination. Together, these findings point to the potential significance of alternative splicing in regulating male development in southern flounder. Further exploration of these genes and their functional implications will deepen our understanding of the role alternative splicing plays in sex differentiation and development.

While alternative splicing genes are well characterized in a range of species, the function of the last putative sex gene we identified, *Groucho* protein, is less clear. *Groucho* acts as a transcriptional corepressor of genes involved in *Drosophila melanogaster* sex determination, and embryos lacking *Groucho* showed defects in sex determination and development [48–49]. To date, *Groucho* family proteins have not been studied in fishes, and further research is needed to determine the role of this marker in male southern flounder development. The discovery of putative sex-related genes, particularly those associated with m6a methylation and alternative splicing, suggests a potential role for epigenetic regulation in sex pathways among northern and southern males.

There was also significantly higher expression of transient receptor potential cation channel transcripts in females. This could be of interest with respect to TSD as this group of proteins serve as environmental sensors in many species. TRPM5 in particular is implicated in temperature sensitivity in at least some contexts [50] and it could therefore be useful to determine which cell types are expressing this protein in developing gonads of paralichthyid flounders.

## Conclusion

Southern flounder exhibit temperature-dependent sex determination (TSD), where genetically XX females develop into phenotypic males when exposed to either lower or higher water temperatures during a critical period in juvenile development. Sex determination in fish is highly dynamic, and, as of now, the specific genes and pathways involved in TSD in southern flounder remain to be more fully characterized.

In this study, we produced the first gonadal transcriptome for southern flounder, and the first transcriptome derived from wild-caught southern flounder. This transcriptome serves as an extensive and previously unavailable resource and offers new insights into the identities of putative sex determination and differentiation genes within southern flounder. A key outcome of our analysis was the identification of candidate sex-determining genes. Through a systematic search for known sex-genes and sex-related Gene Ontology (GO) terms among differentially expressed genes in male and female southern flounder, we identified nine sex-associated genes. Notably, five of these genes had not been previously reported as associated with the sex determination and differentiation process in southern flounder. Given that there is no reference genome available for southern flounder, our functional annotation is based on homologous genes in other species. It is essential to acknowledge that the genes discussed here are putative, representing functional candidates that may not be exact matches to the identified homologous genes. However, the functional annotations were consistently derived from multiple databases employed by Trinotate. Therefore, we are confident that the candidate sex genes serve similar functions to their homologous counterparts; however, further research, including the production of a fully annotated reference genome, is needed to conclusively validate the genes discussed here. This study provides a framework for further investigation of the role these genes play in the sex determination process.

By focusing our search on genes known to be associated with sex in other species, we were able to identify candidate genes that may play a role in the sex determination and differentiation process in southern flounder. However, we also identified both additional differentially expressed genes associated with TGF-β signaling as well as a large suite of sex-biased genes that have not been associated with sex in other species. These other genes could play a role in the sex-determination and differentiation process and should be investigated to further assess their function in southern flounder. In addition, we found many genes expressed in one sex and very low levels or not at all in the other. Biomarkers of gonadal sex informed by our general understanding of gonadal differentiation (*cyp19a1a, foxl2, mis/amh*) have proven useful in determining sex in juvenile southern flounder at body sizes before histological sexing of the gonad is possible [8–9]. This has allowed the assessment of habitat effects on sex determination in wild-caught fish from nursery habitats [24]. While these established biomarkers have proven quite useful, the genes identified in this study as showing still more pronounced sex-biased or sex-specific expression could provide potentially more effective and economical biomarkers for sex identification in developing flounder. More effective and economical biomarkers for phenotypic sexing could potentially strongly benefit both field studies and hatchery work aimed at optimizing production of faster growing females.

As climate change continues to increase ocean temperatures and environmental variability generally [51], fish species with temperature-dependent sex determination (TSD) are likely to face skewed sex ratios and reduced spawning stock biomass. Therefore, investigating the genetic effects of temperature on wild populations is becoming increasingly important. The genes identified in this study can be applied to other teleost species, potentially offering further insights into the TSD process. This study enhances our understanding of TSD and sheds light on key genetic players in the sex determination and differentiation process. Ultimately, this information can be used to better understand how changing environments impact environmentally sensitive marine species.

## Acknowledgements

We would like to thank the NC Division of Marine Fisheries, specifically T. Moore, D. Zapf, T. Wadsworth, C. Stewart and M. Hamric, for providing us with flounder samples.

## Ethics Statement

The research was approved and performed in accordance with the relevant guidelines and regulations by the Institutional Animal Care and Use Committees at North Carolina State University (#13-139-0, 17-003-0).

## Author Contributions

Conceptualization: SH, JM, RB, RG, MBR, JG; Data curation: SH, JM, ES; Formal analysis: SH, ES; Funding acquisition: RB, MBR, JG; Methodology: SH, JM, ES, RB, RG, MBR, JG; Project administration: JM, RG, RB, JG; Resources: RG, RB, JG; Writing – original draft: SH; Writing – review & editing: SH, JM, ES, RB, RG, MBR, JG.

## Data Availability

Raw sequence data and *de novo* transcriptome assembly will be available on Dryad upon acceptance. Differentially expressed genes can be found in the supplementary material.

## Supporting Information Captions

**Table S1.** Genes upregulated in northern males (60-69 mm and 70-79 mm) vs. northern females (60-69 mm and 70-79 mm). Significantly upregulated genes (logFC ≥ 2, FDR < 0.01) are highlighted in gray.

**Table S2.** Genes upregulated in northern females (60-69 mm and 70-79 mm) vs. northern males (60-69 mm and 70-79 mm). Significantly upregulated genes (logFC ≥ 2, FDR < 0.01) are highlighted in gray.

**Table S3.** Genes upregulated in southern males (60-69 mm and 70-79 mm) vs. northern females (60-69 mm and 70-79 mm). Significantly upregulated genes (logFC ≥ 2, FDR < 0.01) are highlighted in gray.

**Table S4.** Genes upregulated in northern females (60-69 mm and 70-79 mm) vs. southern males (60-69 mm and 70-79 mm). Significantly upregulated genes (logFC ≥ 2, FDR < 0.01) are highlighted in gray.

**Table S5.** Differentially expressed genes associated with the TGF-B superfamily.

**Table S6.** Genes unregulated in southern males of 60-69 mm TL vs. southern males of 50-59 mm TL. Significantly upregulated genes (logFC ≥ 2, FDR < 0.01) are highlighted in gray.

**Table S7.** Genes unregulated in southern males of 50-59 mm TL vs. southern males of 60-69 mm TL. Significantly upregulated genes (logFC ≥ 2, FDR < 0.01) are highlighted in gray.

**Table S8.** Genes upregulated in northern males of 60-69 mm TL vs. northern males of 50-59 mm TL. Significantly upregulated genes (logFC ≥ 2, FDR < 0.01) are highlighted in gray.

**Table S9.** Genes upregulated in northern males of 50-59 mm TL vs. northern males of 60-69 mm TL. Significantly upregulated genes (logFC ≥ 2, FDR < 0.01) are highlighted in gray.

**Table S10.** Genes significantly upregulated in both North and South males from 50-59 mm, compared to North and South males 60-69 mm.

**Table S11.** Genes significantly upregulated in both North and South males from 60-69 mm, compared to North and South males 50-59 mm.

